# Characterization of *Salix nigra* floral insect community and activity of three native *Andrena* bees

**DOI:** 10.1101/2020.03.23.004309

**Authors:** Sandra J. Simon, Ken Keefover-Ring, Yong-Lak Park, Gina Wimp, Julianne Grady, Stephen P. DiFazio

## Abstract

*Salix nigra* (black willow) is a widespread tree that hosts many species of polylectic hymenopterans and oligolectic bees of the genus *Andrena*. The early flowering time of *S. nigra* makes it an important nutritive resource for arthropods emerging from hibernation. However, since *S. nigra* is dioecious, not all insect visits will lead to successful pollination. Using both visual observation and pan-trapping we characterized the community of arthropods that visited *S. nigra* flowers and assessed the differences among male and female trees as well as the chemical and visual drivers that influenced community composition across three years. We found that male trees consistently supported higher diversity of insects than female trees and only three insect species, all *Andrena* spp., consistently visited both sexes. Additionally, *A. nigrae*, which was the only insect that occurred more on female than male flowers, correlated strongly to volatile cues. This suggests that cross-pollinators cue into specific aspects of floral scent, but diversity of floral visitors is driven strongly by visual cues. Through time the floral activity of two *Andrena* species remained stable, but *A. nigrae* visited less in 2017 when flowers bloomed earlier than other years. When native bee emergence does not synchronize with bloom, activity appears to be greatly diminished which could threaten species that only subsist on a single host. Despite the community diversity of *S. nigra* flowers, its productivity depends on a small fraction of species that are not threatened by competition, but rather rapidly changing conditions that lead to host-insect asynchrony.

## Introduction

Early spring emergence of flowers is extremely important in supplying nutritive rewards, such as pollen and nectar, for many native arthropods, while the host plant benefits with an increased chance of successful sexual reproduction. The most common cross-pollinators in agricultural systems are often made up of arthropods that collect pollen from many unrelated hosts such as flies belonging to the family Syrphidae, eusocial bee species such as honey bees (*Apis mellifera* L.), and polylectic solitary bees (Ostaff et al. 2015). Lack of discrimination among hosts allows these arthropod groups to more flexibly collect resources for survival and population growth.

Conversely, oligolectic solitary bees, which only collect pollen from either related plant species or a single species, rely heavily on predictable timing of available native floral resources (Danforth 2007; Straka et al. 2014). Upon emergence from nests, oligolectic bees must locate flowers, feed, breed, build new nests, lay eggs, and collect resources to provide their larvae with food provisioned for development throughout the remainder of the year, all within the bloom time of their specific host (Stevens 1949; Linsley 1958; Danforth 2007). In addition to being a valuable resource for early emerging generalist floral insects, willow species belonging to the genus *Salix* are the primary hosts of many oligolectic bees, especially those belonging to one of the largest bee genera, *Andrena* (Stevens 1949; Ostaff et al. 2015).

*Salix* encompasses between 300-400 species of shrubs and trees with a dual pollination system which occurs via wind (anemophilous), insects (entomophilous), or both (Tamura and Kudo 2000; Karrenberg et al. 2002; Argus 2011). *Salix* biology creates a unique environment for insect reward collection due to its dioecious nature. In order for sexual reproduction to occur, insects must locate male plants to collect pollen and then carry it to a separate female plant in the population (Dötterl et al. 2014). Host location typically occurs through a combination of visual, olfactory and reward cues. *Salix* species have a non-showy inflorescence arranged in a catkin form where male flowers are often yellow and female flowers tend to be green (Karrenberg et al. 2002; Füssel et al. 2007). Additionally, *Salix* species emit a complex mixture of volatile organic compounds that are important as olfactory signals to insects, and both male and female plants offer nectar rewards (Tollsten and Knudsen 1992; Füssel et al. 2007). However, upon locating a host plant, some insects may rob flowers of their resources and not carry pollen between male and female individuals (Galen and Butchart 2003).

*Salix nigra*, known by its common name black willow, is a tree-form, entomophilous willow that grows throughout the Eastern United States north to Maine, west to North Dakota and south to Georgia (Burns and Honkala 1990). The extensive range and productivity of *S. nigra* as well as its early bloom, typically February in its southern range through late June in more northern states, makes it an ideal resource for early emerging insects (Burns and Honkala 1990; Ostaff et al. 2015). Studying the mechanisms that *S. nigra* employs to attract insects as well as the influence of sex of tree on floral insect community through time is important in determining the competition for and potential stability of catkin resources, native oligolectic bee activity, and *S. nigra* reproductive success.

The goal of this study was to characterize the community of insects that visit *S. nigra* catkins and examine how the total floral community responded to tree sex, geographic position, volatile organic compound (VOC) profiles, and secondary metabolites in catkins and leaves. For comparison, we also evaluated the community of floral insects captured using visual survey techniques and pan-traps placed in tree canopies. Finally, we examined the effect of tree sex, VOCs, and survey year on the activity of three native *Andrena* bee species, including the willow oligolectic bees *A. macoupinense* and *A. nigrae*.

## Methods

### Population and site description

The target population of *S. nigra* was located in the West Virginia University Core Arboretum in Morgantown, West Virginia. The Core Arboretum is an old growth forest that contains 91 acres of native shrubs, trees, and herbaceous plants. It is located on a hillside that stretches between Monongahela Boulevard and the Monongahela River and contains riparian and floodplain sites with a small grove of *S. nigra* (39.6462° N, 79.9811° W). The population contained thirty-two trees of which twelve were identified as female and twenty were male (Figure 1).

**Figure 1.**
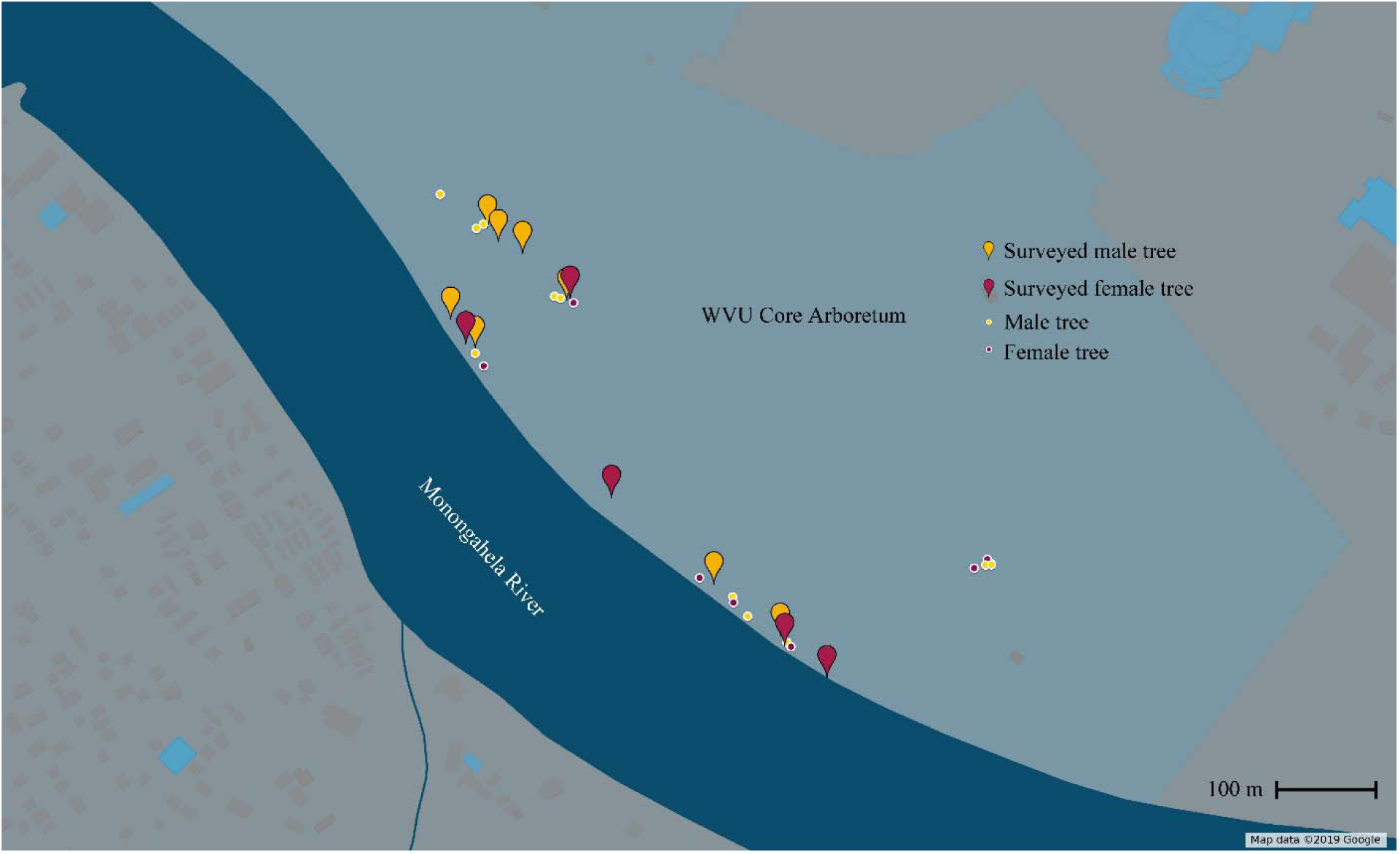
Map of all thirty *Salix nigra* tree locations in WVU Core Arboretum. Pin shapes indicate tree selected for floral insect community survey. Points indicate additional individuals in the Arboretum.

### Visual survey technique

Visual surveys were performed in April through early May in years 2017, 2018 and 2019 on sunny days with minimal wind to help increase observation of small floral visitors. Visual observations were chosen as the survey method for this species due to the brittleness of the base of short shoots, which prevents the use of sweep nets (Beismann et al. 2000). To equalize observations among trees, survey branches were flagged containing approximately 300 catkin flowers for each individual tree. Trees were visually observed for 16 minutes per survey throughout their bloom (∼ 2 weeks) and order of surveyed trees was randomized to account for time of day for two female and four male trees. Insect specimens were carefully hand collected from trees throughout the survey time for family/species level identification. Additionally, survey month and day were recorded for each observation to use as a covariate in models analyzing individual insect activities as well as flower phenology. Finally, early season herbivore activity was collected in 2019 at the end date of tree flowering by counting number of herbivore occurrences on visual survey branches.

### Canopy pan-trap technique

In the year 2019, pan-traps were constructed by painting three-ounce plastic cups with either fluorescent blue, fluorescent yellow, or white paint (Guerra Paint and Pigment, New York). Fluorescent paints were a mixture of 16 ounces of the fluorescent dispersion to 1 gallon of silica flat paint. Three cups, one of each color, were affixed with velcro to a bucket lid as a platform to be raised into tree canopies below flowering branches (Supplemental Figure 1) on five female and six male trees. A soapy trap solution was prepared by addition of approximately 5mL of organic unscented dish soap into one gallon of water. Cups were filled ¾ of the way full with soap solution. Traps were raised to the bottom of each tree canopies (2-12 meters; average 4 meters) at 9:00 am in the morning and emptied daily at 5:00 pm. Visual surveys were performed on the same days. Captured insects from each cup were transferred to separate vials containing 70% ethanol for later identification.

### Insect identification

Abundant insects were identified to species level while rare insects were identified to family. Native bee species identifications were validated by an expert (Sam Droege, Patuxent Wildlife Research Center, personal communication). Vouchers of collected insect specimens were submitted to the West Virginia University Entomology Collection.

### Flower volatile and tissue collection

Dormant branches were collected from the field in early spring for 6 female and 9 male *Salix nigra* trees along the river. Branches were allowed to root and flower in the Biology department greenhouse in buckets of water. Volatile organic compounds (VOCs) were collected using the dynamic headspace method (Keefover-Ring 2013). An oven bag was placed over the flowering branches and bags were secured with thin gauge wire around cotton pads that had been wrapped near the base of the stems. At the top of each bag, polytetrafluoroethylene (PTFE) ports were fixed and connected to chemical traps consisting of 65 mm long and 3 mm internal diameter glass tubes packed with 20 mg of Super Q adsorbent (80/100 mesh size, DVB/ ethylvinylbenzene polymer (Alltech Associates Inc., Deerfield, IL, USA). The traps were connected to calibrated flow meters (Aalborg Instruments & Controls, Inc., Orangeburg, New York, USA) and air was pulled through at a flow rate of 200 ml min^-1^ with an AirLite pump (SKC Inc., Eighty Four, PA) modified to run with a 6 V battery. Controls consisted of oven bags with an identical setup, but without an enclosed branch. Volatiles were collected for a three-hour time period with flow meter maintaining a flow rate of 200 ml min^-1^. At the conclusion of each sampling period, chemical traps were rinsed with 150 µl n-hexane (GC2, Honeywell Burdick & Jackson, Morristown, NJ, USA) into GC vials with PTFE lined screw caps. All catkins were counted in each bag, and maturation stage was noted. All catkins were subsequently lyophilized, and the dry weight obtained of mature and immature (pre-anthesis) catkins separately. Prior to GC-MS analysis, 40 µl of each sample was combined with 2 µl of an internal standard solution (m-xylene in n-hexane). Collected floral volatile organic compounds (VOCs) were analyzed by gas chromatography (GC) with mass spectrometry (MS) detection.

### VOC and metabolite characterization

Floral VOC samples were analyzed with a Thermo Trace 1310 GC coupled to a Thermo ISQ MS with electron ionization (EI) at 70.0 eV at 250 °C, using helium as the carrier gas at 1.0 ml min−1 with the injector temperature set at 250 °C. Oven conditions included an initial temperature of 40 °C followed by an immediate ramp of 3 °C min^−1^ to 200 °C. Available standards, 1 μl of samples, and a continuous series of n-alkanes (C8–C20; Sigma-Aldrich) were injected in the split mode onto a TR-5MS capillary column (30 m × 0.25 mm I.D., film thickness 0.25 μm; Thermo Fisher Scientific). Compounds were identified with retention time matches to pure standards, mass spectra, and/or linear retention indexes calculated with the alkane series (Adams 2007; NIST 2008; El-Sayed 2013). Standard curves of available compounds were used to calculate final VOC results, which were expressed as ng compound g^-1^ DW hr^-1^.

### Sample collection for chemical characterization

Catkins and leaves collected from 24 trees (14 males and 10 females) in the field and in the greenhouse were characterized for five different metabolites; salicin, isosalicin, salicortin, tremuloidin, and termulacin as well as total metabolites. Leaves were flash frozen in the field and later shipped on dry ice to the University of Wisconsin-Madison, WI. The flash frozen catkins and leaves were lyophilized, counted (catkins) and weighed (catkins and leaves), and then ground with steel balls in plastic scintillation vials in a ball mill. Accurately weighed portions (∼15 mg) of powered leaf tissue were extracted with cold (4° C) methanol (1.00 ml) containing salicylic acid-d6 (Sigma-Aldrich, St. Louis, MO, USA) as an internal standard with sonication in an ice bath (15 min) and then centrifugation to obtain a clear supernatant for analysis.

### Chemical analyses

2 μL of all standards and samples solutions were injected onto the UHPLC and separated peaks with a Waters Acquity CSH C-18 column (2.1 x 100 mm, 1.7 μm) at 40°C with a flow rate of 0.5 mL min-1, using a gradient of water (solvent A) and acetonitrile (solvent B), both containing 0.1% formic acid. The PDA was configured to scan from 210-400 nm, with 1.2-nm resolution and a sampling rate of 20 points·s-1. The MS operating parameters for were as follows: cone potential, 30 V; capillary potential, 2500 V; extractor potential, 3 V; RF lens potential, 0.1 V; source temperature, 120 °C; desolvation temperature, 250 °C; desolvation gas flow, 500 L h-1; cone gas flow, 10 L h-1; infusion rate, 5 μL min-1; dwell time, 0.025 s.

Standard curves of methanol solutions, also containing the salicylic acid-d6 internal standard, of various purified compounds were used to calculate the concentrations in the extracted leaves, which were then normalized by sample dry weight and expressed as mg compound g-1. Commercially available standards of salicin (Sigma-Aldrich), and salicortin, HCH-salicortin, tremuloidin, and tremulacin were used that had been previously isolated and purified from aspen foliage (Lindroth et al. 1986).

### Statistical analyses

#### Floral insect community and sex effect on composition

To determine if there were any sex differences in multivariate floral visitor community the R package vegan (Oksanen et al. 2019) was used ordinate the data using non-metric multidimensional scaling (NMDS) using Bray-Curtis dissimilarities. Differences in floral visitor communities among male and female flowers were determined using Analysis of Similarity (ANOSIM), and significance was determined using 999 permutations to determine whether group assignments were significantly different from those generated by chance. Four survey types were tested to determine the stability of sex effects on floral community composition, including: 1) 2019 visual observations, 2) 2019 pan-traps, 3) 2019 total insect community across survey types and 4) visually observed communities through time from 2017-2019. Coordinates for dependent variables were extracted from the NMDS configuration, and depicted in plots using a rescaled font to represent the three dimensional projection. This was accomplished by rescaling all axis scores to a zero origin and scaling the font size relative to the product of the three rescaled axis scores.

#### Sex differences in tree chemistry

NMDS and ANOSIM were also utilized to determine if there were any sex differences in multivariate VOC and metabolite compositions by testing point grouping by sex of tree. Total monoterpenes, sesquiterpenes, VOC emissions, and metabolites were tested using a one-way analysis of variance (ANOVA) in SAS software version 9.4 to determine if there were any differences among sexes.

#### Tree chemistry and insect community relationship

Pairwise geographic distances were calculated among trees from GPS coordinates. VOC production per catkin was scaled to branch level production using average catkin counts per branch for each individual tree to correlate to insect activity. Bray-Curtis dissimilarity matrices were generated for all datasets including floral community, early season herbivore community, pairwise geographic distances, VOCs, and catkin/leaf metabolites. A Mantel test was utilized to determine whether differences in floral insect community were a function of pairwise geographic distances, or differences in catkin VOCs, catkin metabolites or leaf metabolites. This analysis was also repeated for the tree early season herbivore community and catkin/leaf metabolite dissimilarities.

#### 2019 survey method comparison and floral community

Twelve trees (5 female and 7 male) were surveyed in 2019 with visual and pan-trap techniques in the riparian and floodplain sites in the WVU Core Arboretum for a total of one-hundred and forty observations. NMDS and ANOSIM were utilized to determine the effect of survey type on floral community composition. Visual observations were then merged with pan-trap capture counts for overlapping trees and dates for a final fifty-four observations. Independent numeric variables associated with survey day, including Julian date, military time, and temperature, were correlated with the NMDS configuration using the environmental fit vector analysis in vegan (*envfit* function). Variables that were found to significantly correlate to the community dataset were added as covariates in the nested analysis of covariance (ANCOVA) models.

Insects that were abundant in surveys, including *A. macoupinense, A. morrisonella, Andrena nigrae, Lasioglossum coeruleum*, and parasitic wasps belonging to the *Braconidae* family were extracted from datasets. Additionally, species richness was calculated from the dataset as total number of species to visit each tree, and Shannon-Weaver diversity was calculated using the R package vegan. A nested ANCOVA was used to analyze differences in transformed insect counts, richness, and Shannon-Weaver diversity with the following model:

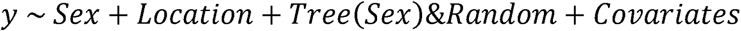

where () indicates the independent variable is nested in another variable.

Additionally, an environmental fit was conducted using the R package vegan with floral volatile compounds which were overlaid on the floral visitor community. The resulting patterns were then evaluated to select specific volatile compounds to look for linear relationships to insect activity using Pearson correlations with Bonferroni p-value corrections to account for multiple testing. Insects chosen to test included *Bombus* sp., *Andrena macoupinense* & *Andrena morrisonella, Andrena nigrae, Miridae, Chalcosyrphus nemorum*, and *Sarcophagidae*. VOCs of interest included acetophenone, cis-b-terpineol-2, ethyl-1-hexanol, germacrene-D, hexenyl-AC, octanal, octen-2-ol, and trans-3-pinanone.

#### Floral community through time

Among all three years, after accounting for mortality and branch loss, a total of six trees along the river overlapped among surveys including two female and four male trees for 77 observation. NMDS and ANOSIM were utilized to determine the effect of survey year on floral insect community composition. Independent numeric variables associated with survey day including year, Julian date, military time, and temperature, and these variables were correlated with the NMDS configuration using the environmental fit vector analysis in vegan (*envfit* function). Variables that were found to significantly correlate to the community dataset were added as additional covariates in all nested ANCOVA models.

Insects that were abundant in surveys, including combined counts of *A. macoupinense, A. morrisonella* and *A. nigrae* were extracted from the community dataset and a nested ANCOVA to analyze differences in transformed insect counts, richness, and Shannon-Weaver diversity with the following model:

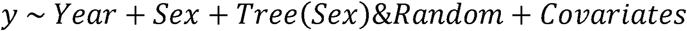

where () indicates the independent variable is nested in another variable.

## Results

### Floral insect community and sex effect on composition

For visual surveys, across all three years of data, 3,160 insects were observed to visit flowers. Of those observations 88.9% were hymenopteran, 9.6% were dipteran, 1.3% were hemipteran and 0.2% were coleopteran. Additionally, of all insects observed, bees belonging to the genus *Andrena* made up 69.4% of the visually surveyed community. NMDS indicated that that the appropriate number of dimensions for 2019 visual surveys, 2019 pan-traps, 2019 total community (Figure 2), and 2017-2019 community (Figure 3) was four (stress = 0.11), three (stress = 0.10), three (stress = 0.12) and four (stress = 0.12) respectively. An ANOSIM indicated that the floral community composition was dependent upon the sex of the tree for all survey types, 2019 total community, and across years (R > 0.1000, p-value < 0.05; Table 1).

**Table 1.**
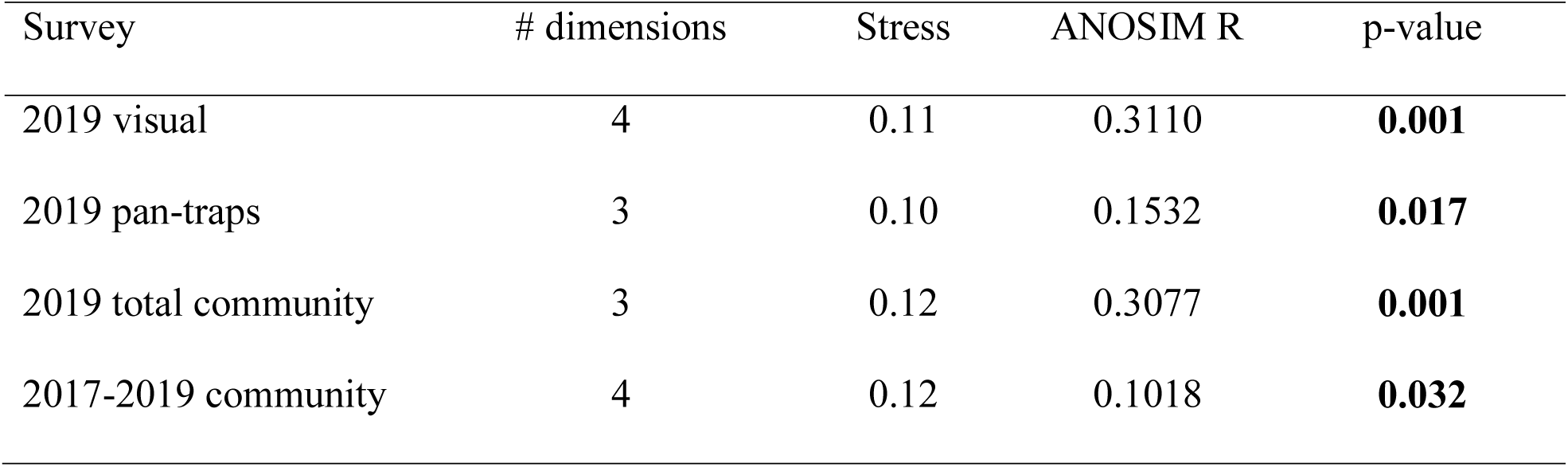
NMDS and ANOSIM (Bray-Curtis dissimilarity) results for multivariate floral visitor composition tested against sex grouping (male vs female). p-values < 0.05 (bolded) indicate that floral visitor composition is more similar within replicate observations of sex group rather than among all observations.

| Survey | # dimensions | Stress | ANOSIM R | p-value |
| --- | --- | --- | --- | --- |
| 2019 visual | 4 | 0.11 | 0.3110 | **0.001** |
| 2019 pan-traps | 3 | 0.10 | 0.1532 | **0.017** |
| 2019 total community | 3 | 0.12 | 0.3077 | **0.001** |
| 2017-2019 community | 4 | 0.12 | 0.1018 | **0.032** |

**Figure 2.**
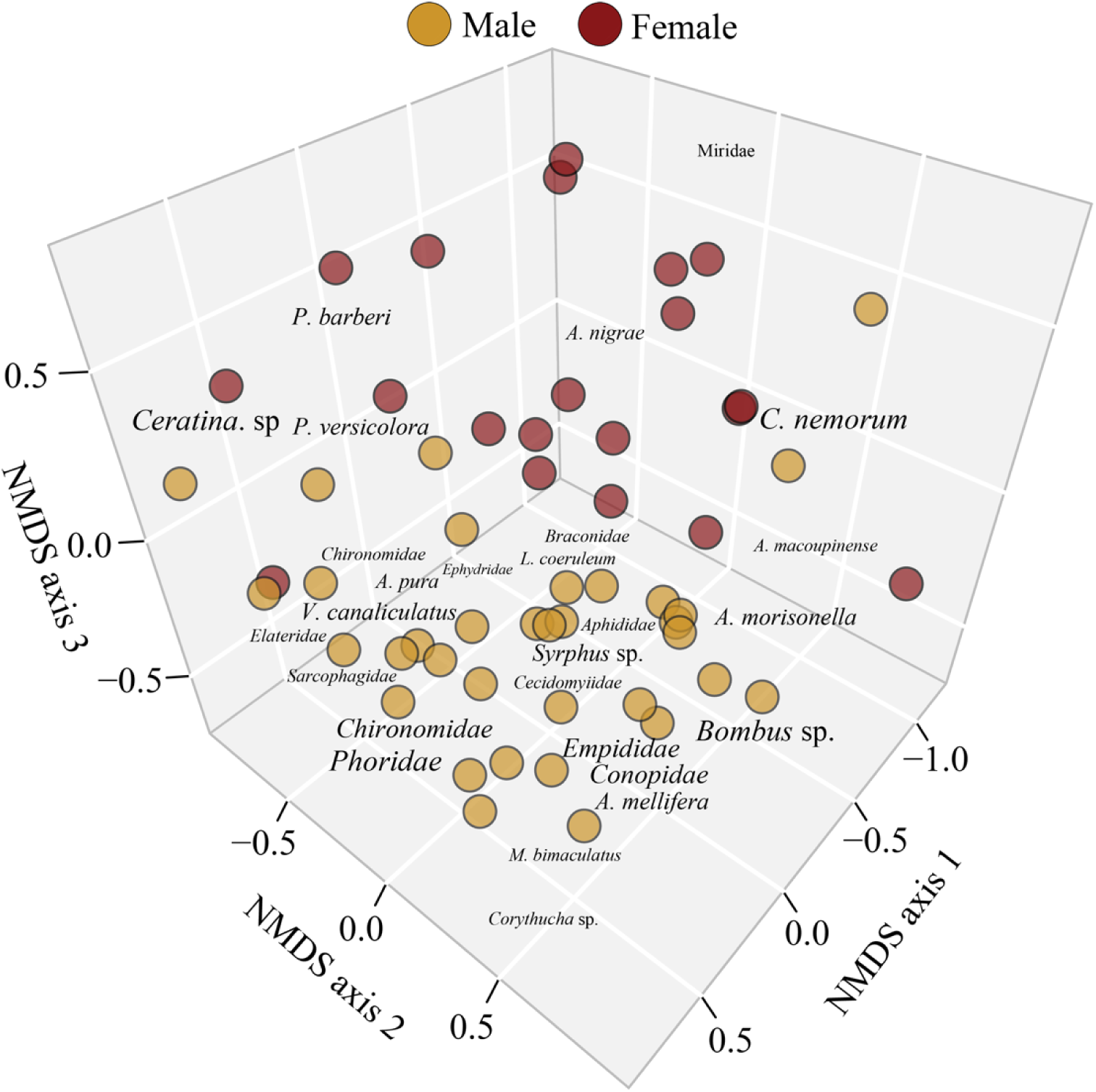
Non-metric multidimensional plot (dimensions = 3; stress = 0.12) of insect floral community with groupings indicated by color for sex of tree (ANOSIM R =0.3077, p-value = 0.001) for 2019 analysis. Font size is scaled to represent three dimensional projection (see Methods).

**Figure 3.**
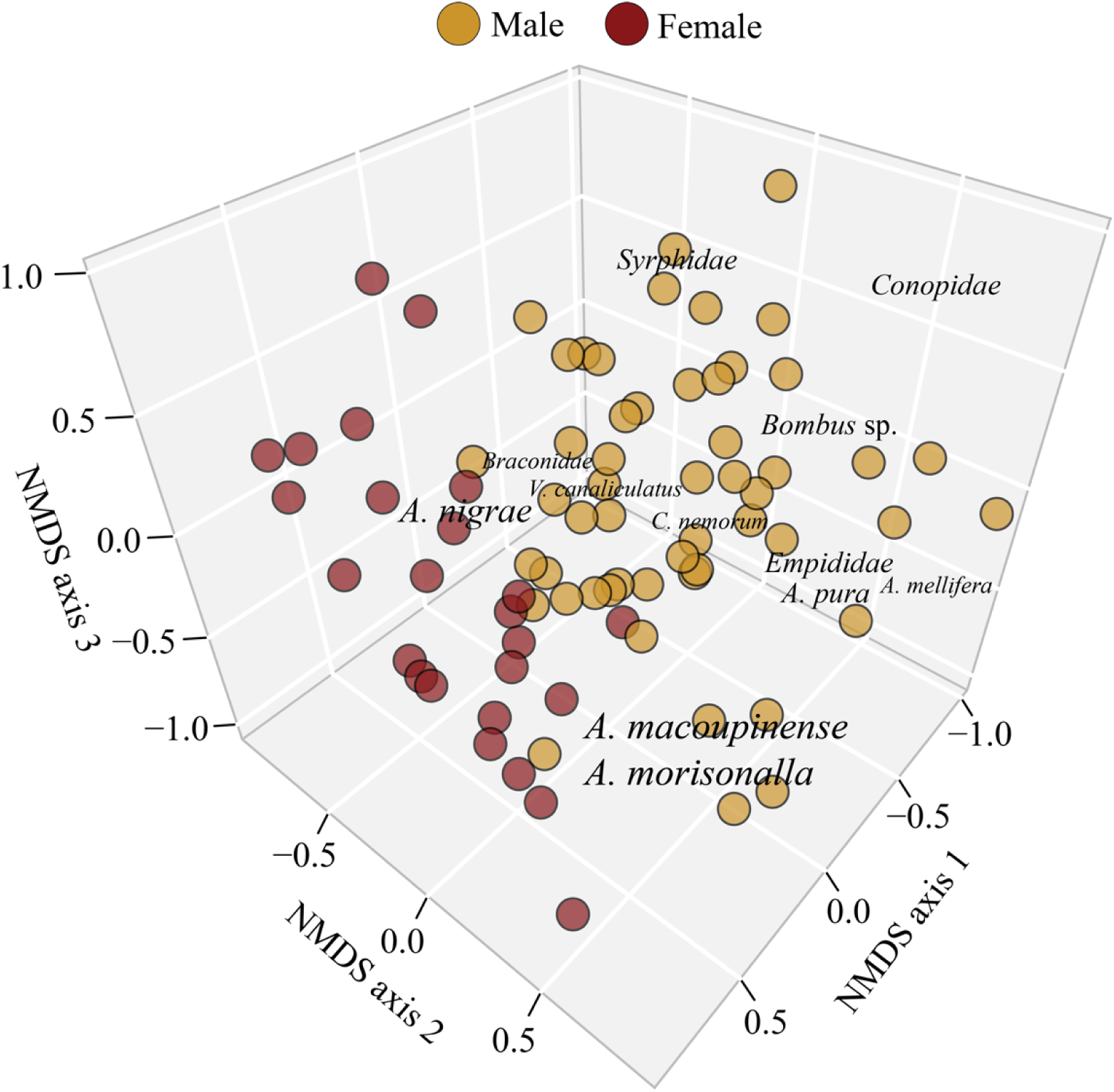
Non-metric multidimensional plot (dimensions = 4; stress = 0.12) of 2017-2019 insect floral community with groupings indicated by color for sex of tree (ANOSIM R = 0.1232, p-value = 0.011).. Font size is scaled to represent three dimensional projection (see Methods).

### Sex differences in tree chemistry

The NMDS of catkin VOCs indicated the appropriate number of dimensions for analysis was three (stress = 0.10) while the number of dimensions for leaf metabolites and catkin metabolites was two (stress = 0.10). An ANOSIM (Table 2) indicated that the catkin VOC composition and leaf metabolite composition were not significantly different between male and female trees (R = -0.1191, p-value = 0.914; R = -0.00471, p-value = 0.449). However, the catkin metabolite composition was different between the sexes (R = 0.1787, p-value = 0.027; Supplemental Figure 2). The total amount of catkin and leaf metabolites did not differ between the sexes (catkins, one-way ANOVA F_1,28_ = 2.741, p-value = 0.109; leaves, F_1,30_ = 1.549, p-value = 0.2229). Furthermore, total monoterpenes (F_1,13_ = 0.02755, p-value = 0.8707), sesquiterpenes (F_1,13_ = 0.2623, p-value = 0.6171), and VOC emissions (F_1,13_ = 0.004865, p-value = 0.9455) were not significantly different between the sexes (Supplemental Figure 3).

**Table 2.** NMDS and ANOSIM (Bray-Curtis dissimilarity) results for multivariate chemistry composition tested against sex grouping (male vs female). p-values < 0.05 (bolded) indicate that chemistry composition is more similar within replicate observations of sex group rather than among all observations.

| Multivariate response | # dimensions | Stress | ANOSIM R | p-value |
| --- | --- | --- | --- | --- |
| VOCs | 3 | 0.10 | -0.1191 | 0.914 |
| Catkin metabolites | 2 | 0.10 | 0.1787 | **0.027** |
| Leaf metabolites | 2 | 0.10 | -0.00471 | 0.449 |

### Tree chemistry and insect community relationship

Mantel tests (Table 3) indicated that there was no relationship between pairwise geographic distances and floral visitor community (r_m_ = 0.026, p-value = 0.450). Similarly, distances between catkin volatile composition, leaf metabolite composition and catkin metabolite composition were not related to differences in floral visitor community (r_m_ = -0.200, p-value = 0.100; r_m_ = 0.002, p-value = 0.480; r_m_ = 0.062, p-value = 0.380, respectively). There was a significant positive relationship between differences in catkin metabolite composition and early season herbivore community (r_m_ = 0.416, p-value = 0.010).

**Table 3.** Mantel test results comparing pairwise Bray-Curtis dissimilarity matrices among insect communities and chemistry composition of flowers and leaves. Bolded p-values and positive r_m_ indicate a significant test, suggesting that similarity in chemistry composition relates to similarity in insect community assemblage.

| Insect matrix | Chemistry matrix | $r_m$ | p-value |
| --- | --- | --- | --- |
| Floral community | Catkin VOCs | -0.300 | 0.07 |
| Floral community | Catkin metabolites | 0.062 | 0.380 |
| Floral community | Leaf metabolites | 0.002 | 0.480 |
| Herbivore community | Catkin metabolites | 0.416 | **0.010** |
| Herbivore community | Leaf metabolites | 0.275 | 0.07 |

### 2019 survey method comparison and floral community

An NMDS with results from both survey methods indicated the appropriate number of dimensions for analysis was three (stress = 0.12; Figure 4a). An ANOSIM indicated that the floral community composition was dependent upon groupings of survey method (R = 0.6656, p-value = 0.001). Pan-traps captured 284 total insects with the majority of insects captured belonging to the orders Diptera (33%) and Coleoptera (41%). There were 1,531 insect observations made during visual surveys with the majority (87%) belonging to the order Hymenoptera (Figure 4b).

**Figure 4.**
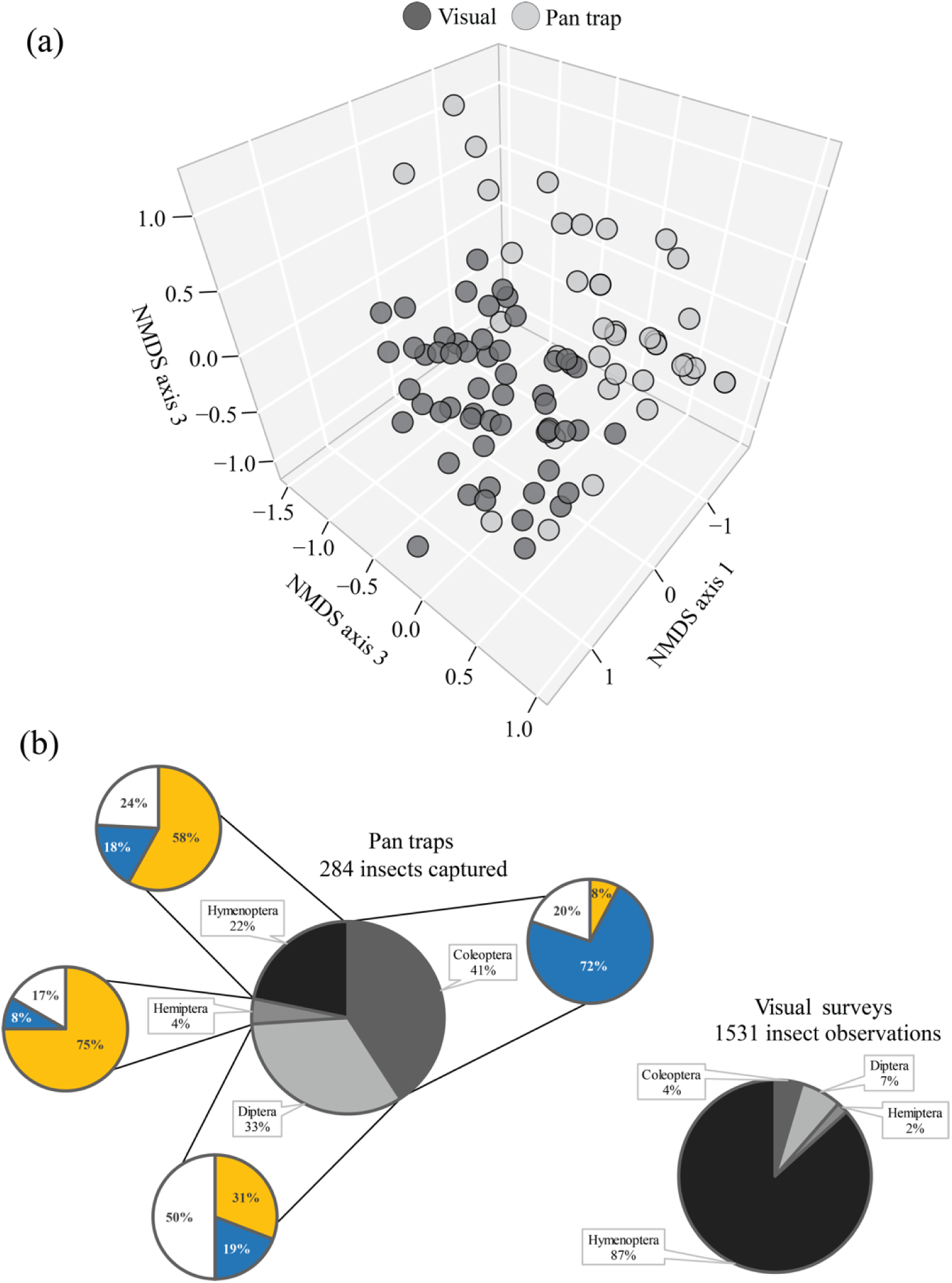
(a) Non-metric multidimensional plot (dimensions = 3; stress = 0.12) of insect floral community with groupings indicated by color for survey method (ANOSIM R = 0.6656, p-value = 0.001). (b) Breakdown of insect capture/observation for two canopy survey methods. Additional pie charts around pan-trap pie chart indicate the percentage of that order captured in different colors of pan-traps.

The NMDS vector analysis indicated that only Julian date was significantly correlated with the NMDS configuration (Vector Max R = 0.3036; p-value = 0.001; Supplemental Table 1). Julian date was then selected for use as a covariate in all nested ANCOVA models. A nested ANCOVA revealed that the occurrence of *A. nigrae* (F_13,40_ = 5.6233; p-value = 0.0310), *A. morrisonella* (F_13,40_ = 7.2722, p-value = 0.0140) and *L. coeruleum* (F_13,40_ = 4.2768, p-value = 0.0500) was dependent on the sex of the tree (Table 4; Figure 4; Supplemental Table 2). Additionally, species richness (F_13,40_ = 17.749; p-value = 0.0004) and Shannon-Weaver diversity (F_13,40_ = 28.936; p-value < 0.0001) also differed between the sexes, with males demonstrating higher values (Table 4; Figure 5; Supplemental Table 3). Finally, the abundance of *A. macoupinense* differed significantly among trees (F_13,40_ = 3.3835, p-value = 0.0030).

**Table 4.** Test of random effects p-values extracted from nested ANCOVA for most abundant floral visitors as well as calculated species richness and Shannon Weaver diversity for 2019 survey analysis. Bolded values indicate that the independent variable had a significant effect on the dependent variable (p-value < 0.05).

|  | *Andrena nigrae* | *Andrena macoupinense* | *Andrena morrisonella* | *Lasioglossum coeruleum* | Species richness | Shannon-Weaver diversity |
| --- | --- | --- | --- | --- | --- | --- |
| Sex | **0.031** | 0.5447 | **0.014** | **0.0500** | **0.0004** | **<0.0001** |
| Location | 0.5645 | 0.3587 | 0.8326 | 0.9577 | 0.9536 | 0.7348 |
| Tree | 0.2046 | **0.003** | 0.4729 | 0.1635 | 0.5944 | 0.6917 |
| Julian date | 0.5645 | 0.6261 | 0.0600 | 0.6141 | 0.5800 | 0.2839 |

**Figure 5.**
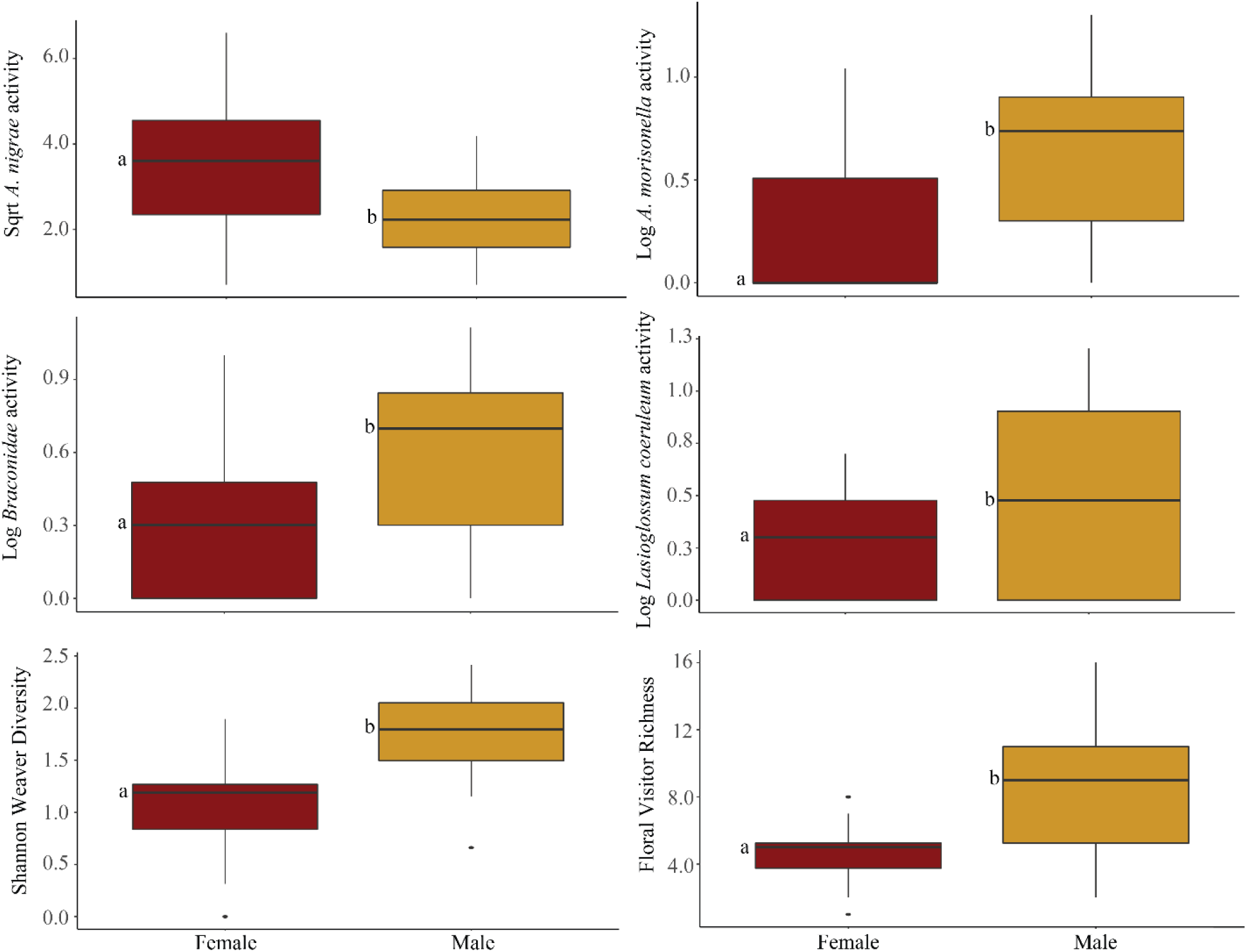
Average activity of most common floral visitors and calculated community metrics from 2019 *S. nigra* surveys. Letters to the left of boxes indicate significantly different means as determined by a Tukey’s HSD (p-value < 0.05).

A significant correlation was found between the abundance of *A. nigrae* and two volatile compounds; acetophenone (ρ *=* 0.9122, p-value = 0.05) and octen-2-ol (ρ *=* 0.9393, p-value = 0.01) (Supplemental Figure 4). Additional VOCs and insect activities chosen to test were not significant (p-value > 0.05).

### Floral community through time

An ANOSIM indicated that the floral community composition differed among years (R = 0.1669, p-value = 0.001). The vector analysis indicated that only Julian date was significantly correlated with the NMDS dimensions (Vector Max R = 0.0802; p-value = 0.047; Supplemental Table 4). A nested ANCOVA (Table 5; Supplemental Figure 5; Supplemental Table 5) showed that average species richness (F_8,68_ = 32.2841, p-value = 0.0046) and Shannon-Weaver diversity (F_8,68_ = 34.6569; p-value = 0.0041) were dependent on the sex of the tree. Additionally, flowering time occurred substantially earlier in 2017 than in the other survey years (Figure 6a) which may be related to the dependence of *A. nigrae* activity on the survey year (F_8,68_ = 5.1049; p-value = 0.0086; Figure 6b).

**Table 5.** Test of random effects p-values extracted from nested ANCOVA for most abundant floral visitors as well as calculated species richness and Shannon Weaver diversity. Bolded values indicate that the independent variable had a significant effect on the dependent variable (p-value < 0.05).

|  | *Andrena nigrae* | Species richness | Shannon-Weaver diversity |
| --- | --- | --- | --- |
| Sex | 0.1475 | **0.0046** | **0.0041** |
| Year | **0.0086** | 0.7303 | 0.9019 |
| Tree | 0.1134 | 0.4356 | 0.2488 |
| Julian date | 0.2171 | 0.2467 | 0.2617 |

**Figure 6.**
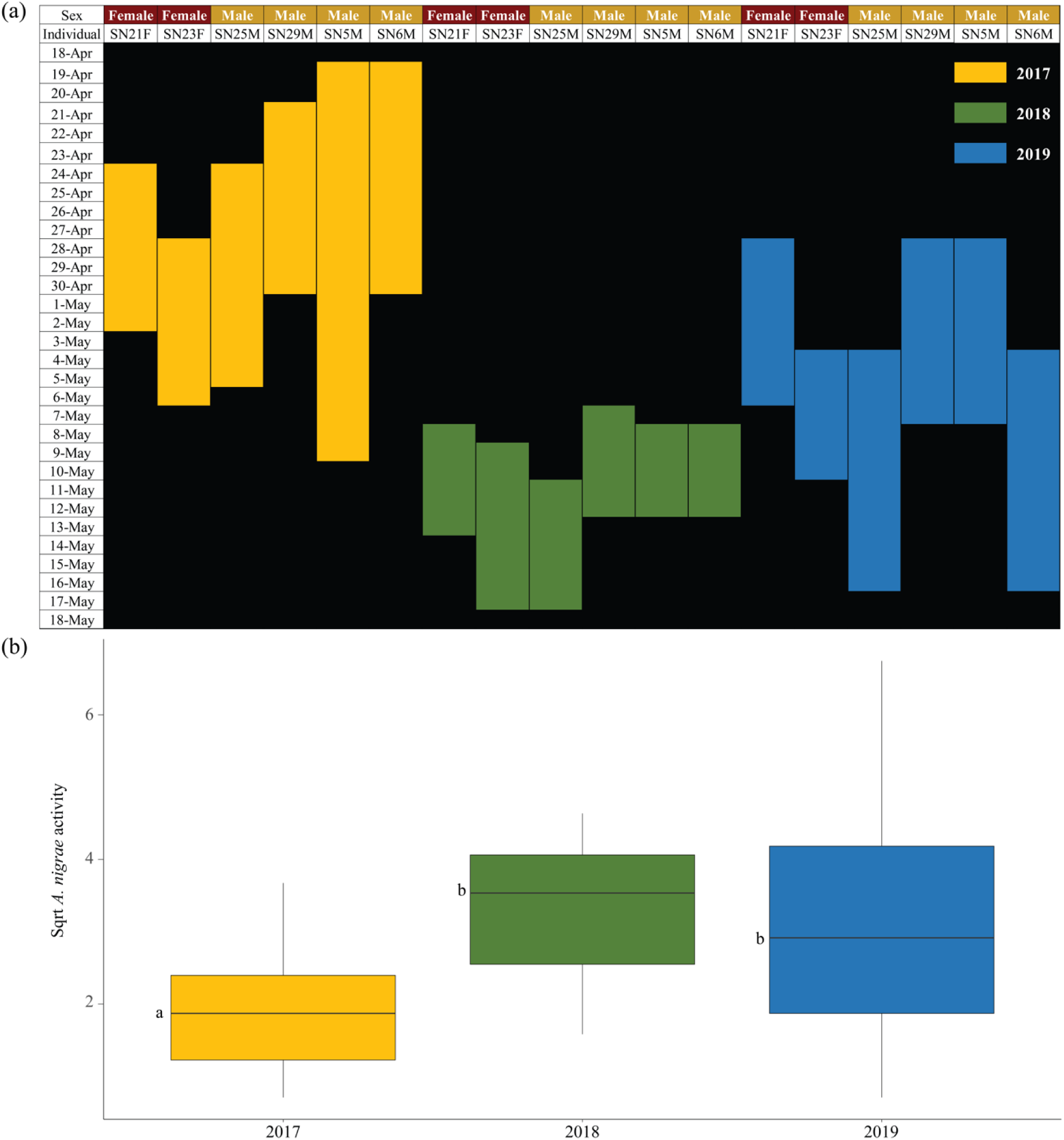
(a) Amount of time and date in each year (2017, 2018, and 2019) in which catkins were actively attractive to insects. End date for male individuals indicates trees have dropped all catkins while female trees no longer have receptive stigmas (brown and shriveled) or active insect in canopies**. (**b) Average count of *A. nigrae* visits to *S. nigra* catkins in survey years 2017, 2018 and 2019 from year analysis model. Letters to the left of boxes indicate significantly different means as determined by a Tukey’s HSD (p-value < 0.05).

## Discussion

The total floral community composition was strongly influenced by the sex of tree across survey techniques and in all years. Male trees consistently attracted a more diverse and unique insect assemblage on their flowers when compared to female trees. However, there was no relationship detected between the sex of the tree and the floral VOC composition indicating that the scent of male and female flowers among trees was not different. Total emission of monoterpenes, sesquiterpenes, and all VOCs were also found to be the same among sexes. Similarly, differences in floral scent and plant tissue defenses among trees were not strong drivers of the community of insects that visited the flowers. Thus, we found no evidence that the influence that sex of tree on floral insect assemblages was due to the effects of VOCs or metabolite composition, though more subtle relationships might be revealed with more intensive sampling.

Our results are unusual given that dioecious plants, in which both sexes offer rewards, are often dimorphic in both scent composition and emission levels (Füssel 2007; Ashman 2009; Okamoto et al. 2013). Nevertheless, this pattern does not always hold true in the *Salix* genus. For example, *S. repens, S. bicolor, S. caprea*, and *S. cinerea* all have the same overall volatile composition among male and female flowers (Tollsten and Knudsen 1992; Füssel 2007). This may reflect differences in the balance between visual and olfactory cues among *Salix* species, though this hypothesis remains to be robustly tested.

Although male and female trees did not differ in VOC and leaf metabolite composition, there was a sex effect on the metabolite composition of catkins. We also found that catkin metabolites exerted a strong influence over herbivore composition, suggesting that floral defenses are utilized differently among male and female trees, in turn attracting unique assemblages of herbivores. In the early season, trees may be at risk of herbivores feeding directly on flowers which act as a large resource sink in plant tissues (Wäckers et al. 2007). Floral larceny is a threat to female reproduction since they must maintain flowers through seed production, and this relationship may be antagonized in a dioecious system where chance of accidental pollination is rare (Maloof and Inouye 2000; Richardson 2004). Thus, it is important that females invest more resources toward defense of catkins, as supported by our finding that female individuals emitted more (3*E*)-Hexenyl acetate and (2*E*)-Hexenyl acetate. Hexenyl acetates have been frequently characterized as common green leaf volatiles emitted upon the crushing of plant tissue (Whitman and Eller 1990; Heil et al. 2008; Wei and Kang 2011) and they appear to also be a common component in floral scent (Tollsten and Knudsen 1992; Kaiser 1994; Messinger 2006). Hexenyl acetates are associated with the attraction of insect parasites which may provide female flowers in *S. nigra* added protection benefits against floral resource theft (Whitman and Eller 1990; Ngumbi et al. 2009).

In our 2019 floral community surveys pan-traps were employed to test differences in surveys techniques and characterize total insect community shifts among male and female trees. We determined that pan-trapping and visual survey techniques captured vastly different samples of the floral visitor community. Pan-traps were very effective at capturing small insects which belonged to Diptera and Coleoptera, explaining much of the disparity between techniques. However, hymenopterans made up 87% of insects counted during visual surveys with 67% of bee observations coming from native *Andrena* spp. Conversely, hymenopterans only made up 22% of insects captured in pan-traps with only 8% of the bees captured belonging to *Andrena* spp. Unscented pan-traps filter the insect community, which were presumably attracted to the flowers on the tree, down to those relying heavily on visual cues (Laubertie et al. 2006; Tuell and Isaacs 2009; Vrdoljak and Samways 2012). The lack of native *Andrena* spp. attraction to pan-traps may indicate that both visual and volatile cues are important in their association rather than only visual cues. Additionally, given that the pan-trap data increased the diversity disparity observed between male and female flowers in 2019, it appears that the initial visual component of male pollen may be influencing the differences in the larger floral community shift observed among male and female trees. This is further supported by the differential rates of capture of the hemipterans and hymenopterans in yellow cups and coleopterans in blue cups.

Surveys completed in 2019 revealed the sex of tree was important for many of the Hymenopteran distributions throughout our site. *A. morisonella* as well as *L. coeruleum* and a parasitoid wasp species in the family Braconidae all showed higher activity on male trees indicating a prioritization of male rewards. *A. macoupinense* was equally observed among male and female trees. Interestingly, *A. nigrae* was more actively visiting female flowers on trees in the study site. Given the non-showy nature of the *Salix* female catkins this suggests that the species may be more tightly coupled to the volatile rather than visual cues in order to locate an appropriate host. In support of this *A. nigrae* was the only insect whose activity was significantly correlated to level of specific floral volatiles. The number of visits of *A. nigrae* increased with increasing levels of both acetophenone and octen-2-ol.

Despite detection of sex differences in activity of native bees in 2019, *Andrena* spp. were stable among male and female flowers through time, indicating that all three species are important in contributing to sexual reproduction of *S. nigra*. Additionally, the total activity of *A. macoupinense* and *A. morisonella* were stable across years, but *A. nigrae* had far fewer catkin visits in 2017 when compared to later years. Adult emergence of *A. macoupinense* has been recorded mid-March through May, *A. nigrae* is often found April through May, and *A. morisonella* is frequent from May to early June (Stevens 1949; Ribble 1967, 1974a, b). Of the three species, *A. nigrae* emergence is more closely coupled to the bloom of local *S. nigra* trees. Additionally *A. nigrae* has been recorded as a primary pollinator of *S. nigra* in two other states, Illinois and North Dakota, indicating that it may be more tightly coupled to the biology of *S. nigra* than other observed native bees in our site (Stevens 1949; Ribble 1967).

When examining the phenology of flowers through time, the bloom time of trees in 2017 occurred earlier than subsequent years due to uncharacteristically warm weather in early March followed by much cooler temperatures through April and early May. The abnormality of temperature may have decoupled the availability of floral resources from local *A. nigrae* population emergence. Additionally, cooler temperatures may have had a negative effect on volatilization of catkin scent leading to the inability of native bees to effectively locate their host. The lack of activity difference detected for *A. macoupinense* and *A. morisonella* may indicate that local populations effectively supplement their diets with resources from additional plant species such as other local *Salix*, including *S. discolor* and *S. bebbiana*, or *Prunus* and *Amelanchier* spp., both of which are native to West Virginia and listed as occasional hosts (Ribble 1974a, b).

While they attract a more diverse community of floral visitors, male individuals of *S. nigra* also overproduce resources (average pollen:ovule ratio ∼1500:1, J. Grady, unpublished data) thus minimizing competition between native pollinators and pollen/nectar thieves. Conversely, female individuals lack strong visual cues and, despite having similar floral volatile composition, appear to have different protective chemistry composition in their catkins allowing them to preserve their rewards directly for the cross-pollinating *Andrena* species. Competition for resources does not seem to be a direct threat to *Andrena* activity or successful sexual reproduction of *S. nigra*. However, we show that one native bee species may be vulnerable when local tree bloom time is decoupled from their seasonal emergence. For most native pollinators and plants, partial asynchrony with bloom time is not a threat if additional hosts are present and insect phenology has time to adjust to permanently altered conditions such as those caused by climate change (Willmer 2012; Bartomeus et al. 2013). For oligolectic species like *A. nigrae* that rarely collect from hosts outside of a single plant group or species, even mild decoupling could have drastic implications for locally adapted bee populations. As a result, plant sexual reproduction will be negatively impacted, especially when the cross-pollinating community is made up of specialists, and accidental pollination becomes rare.

Overall, differences in floral community composition and diversity among male and female trees of *S. nigra* appears to be strongly driven by visual cues. However, the main cross pollinators, *Andrena* spp., rely on both visual and specific volatile cues to locate male individuals and carry pollen to the less showy female trees. Additionally, the interannual variability in flowering time highlights the potential effects of climate on pollinator activity and tree sexual productivity. Inter-annual variability in the *S. nigra* – *Andrena* interaction illustrates that shifting seasonal transitions could detrimentally affect plants that depend on early-emerging arthropods for sexual reproduction as well as the arthropods that depend on resources provided by early-flowering plants.

## Acknowledgements

We thank the following individuals for assistance with fieldwork-Jacob Miller, Brandon Sinn, and Mark Burnham. Zachariah Fowler provided logistical support and access to the WVU Core Arboretum *Andrena* identification was performed by Sam Droege Craig Larcenaire provided advice on pan traps. Funding was provided by the NSF Dimensions of Biodiversity Program (DEB-1542509 to S.D. and DEB-1542479 ti K.K.-R.), and by a fellowship from the Department of Biology of West Virginia University to S.J.S.).

## References

Adams RP (2007) Identification of essential oil components by gas chromatography/mass spectrometry, vol. 456. Allured publishing corporation, Carol Stream, IL

Argus GW (2011) An experimental study of hybridization and pollination in *Salix* (willow). Artic Can J Bot 52:1613–1619. doi: 10.1139/b74-212

Ashman TL (2009) Sniffing out patterns of sexual dimorphism in floral scent. Funct Ecol 23:852–862. doi: 10.1111/j.1365-2435.2009.01590.x

Bartomeus I, Park MG, Gibbs J, et al (2013) Biodiversity ensures plant-pollinator phenological synchrony against climate change. Ecol Lett 16:1331–1338. doi: 10.1111/ele.12170

Beismann H, Wilhelmi H, Baillères H, et al (2000) Brittleness of twig bases in the genus *Salix*: fracture mechanics and ecological relevance. J Exp Bot 51:617–633. doi: 10.1093/jexbot/51.344.617

Burns RM, Honkala BH (1990) Silvics of North America: Volume 2. Hardwoods. United States Dep Agric (USDA), For Serv Agric Handb 654

Danforth B (2007) Bees. Curr Biol 17:156–161. doi: 10.1016/j.cub.2007.01.025

Dötterl S, Glück U, Jürgens A, et al (2014) Floral reward, advertisement and attractiveness to honey bees in dioecious *Salix caprea*. PLoS One 9:e93421. doi: 10.1371/journal.pone.0093421

El-Sayed AM (2013) The pherobase: database of insect pheromones and semiochemicals. http://www.pherobase.com/guide/

Füssel U (2007) Floral scent in *Salix* L. and the role of olfactory and visual cues for pollinator attraction of *Salix caprea* L. PhD dissertation, Department of Biology, Chemistry and Sciences, Bayreuth University, Bayreuth, Germany

Füssel U, Dötterl S, Jürgens A, Aas G (2007) Inter- and intraspecific variation in floral scent in the genus *Salix* and its implication for pollination. J Chem Ecol 33:749–765. doi: 10.1007/s10886-007-9257-6

Galen C, Butchart B (2003) Ants in your plants: effects of nectar-thieves on pollen fertility and seed-siring capacity in the alpine wildflower, Polemonium viscosum. Oikos 101:521–528. doi: 10.1034/j.1600-0706.2003.12144.x

Heil M, Lion U, Boland W (2008) Defense-inducing volatiles: in search of the active motif. J Chem Ecol 34:601–604. doi: 10.1007/s10886-008-9464-9

Kaiser R (1994) Trapping, investigation and reconstitution of flower scents. Springer Netherlands

Karrenberg S, Edwards P, Kollmann J (2002) The life history of Salicaceae living in the active zone of floodplains. Freshw Biol 47:733–748. doi: 10.1046/j.1365-2427.2002.00894.x

Keefover-Ring K (2013) Making scents of defense: do fecal shields and herbivore-caused volatiles match host plant chemical profiles? Chemoecology 23:1–11. doi: 10.1007/s00049-012-0117-7

Laubertie EA, Wratten SD, Sedcole JR (2006) The role of odour and visual cues in the pan-trap catching of hoverflies (Diptera: Syrphidae). Ann Appl Biol 148:173–178. doi: 10.1111/j.1744-7348.2006.00046.x

Lindroth RL, Scriber JM, Hsia MTS (1986) Differential responses of tiger swallowtail subspecies to secondary metabolites from tulip tree and quaking aspen. Oecologia 70:13–19. doi: 10.1007/BF00377106

Linsley EG (1958) The ecology of solitary bees. Hilgardia 27:543–599. doi: 10.3733/hilg.v27n19p543

Maloof JE, Inouye DW (2000) Are nectar robbers cheaters or mutualists? Ecology 81:2651–2661.

Messinger OJ (2006) The role of visual and olfactory cues in host recognition for the specialist bee genus Diadasia, and implications for the evolution of host choice. PhD dissertation, Department of Entomology, Southern Illinois University at Carbondale

Ngumbi E, Chen L, Fadamiro HY (2009) Comparative GC-ead responses doi:10.1890/0012-9658(2000)081[2651:ANRCOM]2.0.CO;2of a specialist (Microplitis croceipes) and a generalist (Cotesia marginiventris) parasitoid to cotton volatiles induced by two caterpillar species. J Chem Ecol 35:1009–1020. doi: 10.1007/s10886-009-9700-y

NIST (2008) National institute of standards and technology mass spectral library. National Institute of Standards and Technology, US Department of Commerce, Washington, D.C.

Okamoto T, Kawakita A, Goto R, et al (2013) Active pollination favours sexual dimorphism in floral scent. Proc R Soc B Biol Sci 280:. doi: 10.1098/rspb.2013.2280

Oksanen J, Blanchet FG, Friendly M, et al (2019) vegan: community ecology package

Ostaff DP, Mosseler A, Johns RC, et al (2015) Willows (*Salix* spp.) as pollen and nectar sources for sustaining fruit and berry pollinating insects. Can J Plant Sci 95:505–516. doi: 10.4141/CJPS-2014-339

Ribble DW (1974a) A Revision of the Bees of the Genus Andrena of the Western Hemisphere Subgenus Scaphandrena. Am. Entomol. Soc. 100:101–189

Ribble DW (1974b) A Revision of the Bees of the Genus Andrena of the Western Hemisphere Subgenus *Scaphandrena*. Am Entomol Soc 100:101–189

Ribble DW (1967) Revisions of two subgenera of *Andrena: Micrandrena* (Ashmead) and *Derandrena*. PhD dissertation, Department of Entomology, University of Nebraska

Richardson SC (2004) Are nectar-robbers mutualists or antagonists? Oecologia 139:246–254. doi: 10.1007/s00442-004-1504-8

Stevens OA (1949) Native bees. Exp Stn Bimon Bull 12:90–98

Straka J, Rina Cern K, Mach L, et al (2014) Life span in the wild: the role of activity and climate in natural populations of bees. Funct Ecol 28:1235–1244. doi: 10.1111/1365-2435.12261

Tamura S, Kudo G (2000) Wind pollination and insect pollination of two temperate willow species, *Salix miyabeana* and *Salix sachalinensis*. Plant Ecol 147:185–192. doi: 10.1023/A:1009870521175

Tollsten L, Knudsen JT (1992) Floral scent in dioecious *Salix* (Salicaceae)-a cue determining pollination system? Plant Syst Evol 182:229–237. doi: 10.1007/BF00939189

Tuell JK, Isaacs R (2009) Elevated pan traps to monitor bees in flowering crop canopies. Entomol Exp Appl 131:93–98. doi: 10.1111/j.1570-7458.2009.00826.x

Vrdoljak SM, Samways MJ (2012) Optimising coloured pan traps to survey flower visiting insects. J Insect Conserv 16:345–354. doi: 10.1007/s10841-011-9420-9

Wäckers FL, Romeis J, van Rijn P (2007) Nectar and pollen feeding by insect herbivores and implications for multitrophic interactions. Annu Rev Entomol 52:301–323. doi: 10.1146/annurev.ento.52.110405.091352

Wei J, Kang L (2011) Roles of (Z)-3-hexenol in plant-insect interactions. Plant Signal Behav 6:369–371. doi: 10.4161/psb.6.3.14452

Whitman DW, Eller FJ (1990) Parasitic wasps orient to green leaf volatiles. Chemoecology 1:69–76. doi: 10.1007/BF01325231

Willmer P (2012) Ecology: Pollinator–Plant Synchrony Tested by Climate Change. Curr Biol 22:R131–R132. doi: 10.1016/j.cub.2012.01.009

