## Supplemental Figures and Tables for "Characterization of *Salix nigra* floral insect community and activity of three native *Andrena* bees"

**Supplemental Table 1.** Environmental fit results for 2019 floral community NMDS analysis. Bolded p-values (<0.05) indicate a significant correlation of the independent variable with the NMDS configuration.

| Independent variable | Vector Max R | p-value |
|---|---|---|
| Julian date | 0.3036 | **0.001** |
| Temperature | 0.0582 | 0.241 |
| Military time | 0.0447 | 0.308 |

**Supplemental Table 2.** Nested ANCOVA model results for 2019 analysis of individual native bee abundances. Bolded p-values (<0.05) indicate significant model effects.
y ~ Sex + Location + Tree(Sex)&Random + Julian date

| Dependent variable: *Andrena nigrae* | | | | | |
|---|---|---|---|---|---|
| Source | Degrees of freedom | Sum of squares | Mean square | F-Ratio | p-value |
| Model | 13 | 43.48185 | 3.34476 | 2.4947 | **0.0134** |
| Error | 40 | 53.62887 | 1.34072 | | |
| C. Total | 53 | 97.11072 | | | |

| Dependent variable: *Andrena macoupinense* | | | | | |
|---|---|---|---|---|---|
| Source | Degrees of freedom | Sum of squares | Mean square | F-Ratio | p-value |
| Model | 13 | 5.604838 | 0.431141 | 3.9674 | **0.0004** |
| Error | 40 | 4.34681 | 0.10867 | | |
| C. Total | 53 | 9.951648 | | | |

| Dependent variable: *Andrena morrisonella* | | | | | |
|---|---|---|---|---|---|
| Source | Degrees of freedom | Sum of squares | Mean square | F-Ratio | p-value |
| Model | 13 | 4.347317 | 0.334409 | 2.9616 | **0.0042** |
| Error | 40 | 4.516573 | 0.112914 | | |
| C. Total | 53 | 8.86389 | | | |

| Dependent variable: *Lasioglossum coeruleum* | | | | | |
|---|---|---|---|---|---|
| Source | Degrees of freedom | Sum of squares | Mean square | F-Ratio | p-value |
| Model | 13 | 3.101719 | 0.238594 | 2.2816 | **0.023** |
| Error | 40 | 4.182974 | 0.104574 | | |
| C. Total | 53 | 7.284693 | | | |

**Supplemental Table 3.** Nested ANCOVA model results for 2019 analysis of calculated community metrics. Bolded p-values (<0.05) indicate significant model effects.
y ~ Sex + Location + Tree(Sex)&Random + Julian date

| Dependent variable: Species richness | | | | | |
|---|---|---|---|---|---|
| Source | Degrees of freedom | Sum of squares | Mean square | F-Ratio | p-value |
| Model | 13 | 307.5722 | 23.6594 | 3.3871 | **0.0015** |
| Error | 40 | 279.4093 | 6.9852 | | |
| C. Total | 53 | 586.9815 | | | |

| Dependent variable: Shannon-Weaver diversity | | | | | |
|---|---|---|---|---|---|
| Source | Degrees of freedom | Sum of squares | Mean square | F-Ratio | p-value |
| Model | 13 | 9.700905 | 0.746223 | 5.2588 | **<0.0001** |
| Error | 40 | 5.675971 | 0.141899 | | |
| C. Total | 53 | 15.37688 | | | |

**Supplemental Table 4.** Environmental fit results for floral community NMDS analysis across years (2017-2019). Bolded p-values (<0.05) indicate a significant correlation of the independent variable with the NMDS configuration.

| Independent variable | $r^2$ | p-value |
|---|---|---|
| Year | 0.0802 | **0.047** |
| Julian date | 0.0815 | **0.046** |
| Temperature | 0.0491 | 0.18 |

**Supplemental Table 5.** Nested ANCOVA model results for year analysis. Bolded p-values (<0.05) indicate significant model effects.
y ~ Sex + Year + Tree(Sex)&Random + Julian date

| Dependent variable: *Andrena macoupinense* & *Andrena morrisonella* | | | | | |
|---|---|---|---|---|---|
| Source | Degrees of freedom | Sum of squares | Mean square | F-Ratio | p-value |
| Model | 8 | 27.17312 | 3.39664 | 1.6702 | 0.1218 |
| Error | 68 | 138.2903 | 2.03368 | | |
| C. Total | 76 | 165.4635 | | | |

| Dependent variable: *Andrena nigrae* | | | | | |
|---|---|---|---|---|---|
| Source | Degrees of freedom | Sum of squares | Mean square | F-Ratio | p-value |
| Model | 8 | 46.10628 | 5.76328 | 4.9103 | **<0.0001** |
| Error | 68 | 79.81222 | 1.17371 | | |
| C. Total | 76 | 125.9185 | | | |

| Dependent variable: Species richness | | | | | |
|---|---|---|---|---|---|
| Source | Degrees of freedom | Sum of squares | Mean square | F-Ratio | p-value |
| Model | 8 | 82.47375 | 10.3092 | 4.8422 | **<0.0001** |
| Error | 68 | 144.773 | 2.129 | | |
| C. Total | 76 | 227.2468 | | | |

| Dependent variable: Shannon-Weaver diversity | | | | | |
|---|---|---|---|---|---|
| Source | Degrees of freedom | Sum of squares | Mean square | F-Ratio | p-value |
| Model | 8 | 5.76738 | 0.720923 | 7.3942 | **<0.0001** |
| Error | 68 | 6.62987 | 0.097498 | | |
| C. Total | 76 | 12.39725 | | | |

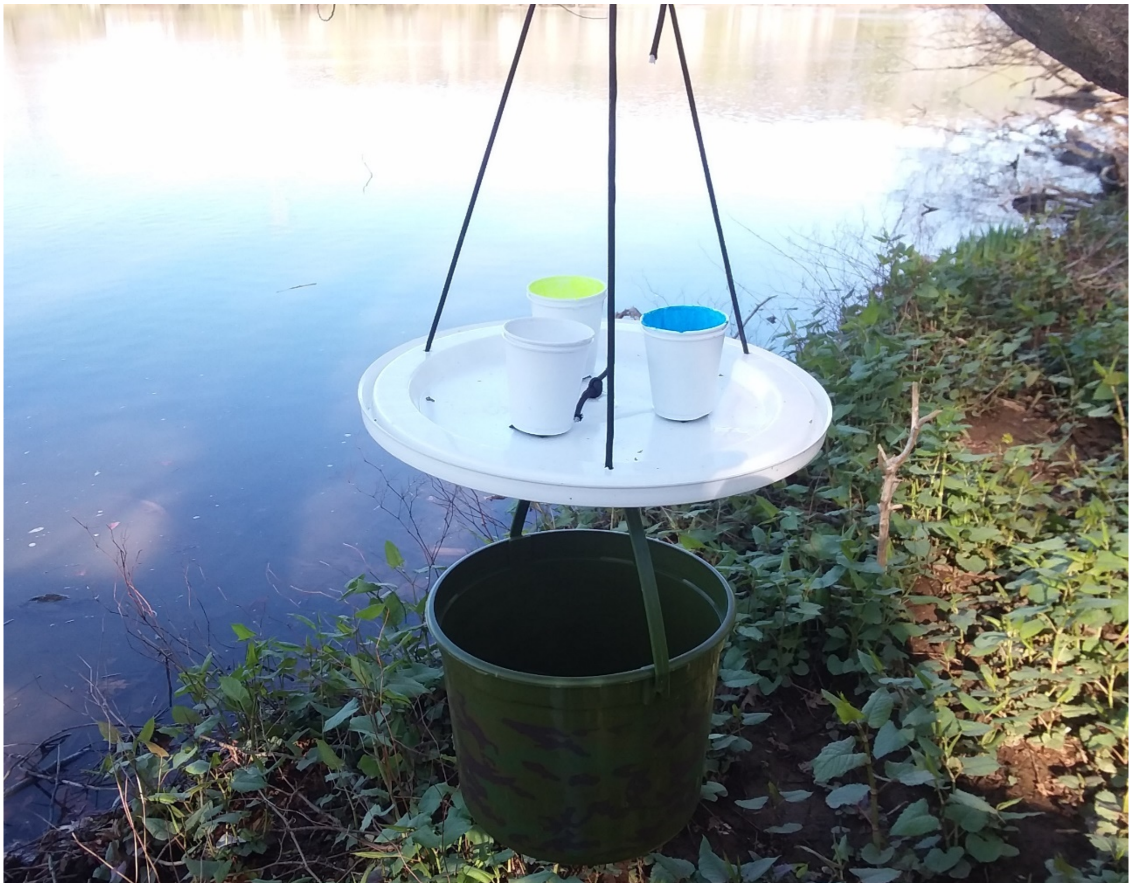

**Supplemental Figure 1.** Tree canopy pan-trap design. Camouflaged buckets were added to hang below river traps to add weight, preventing winds from jostling traps.

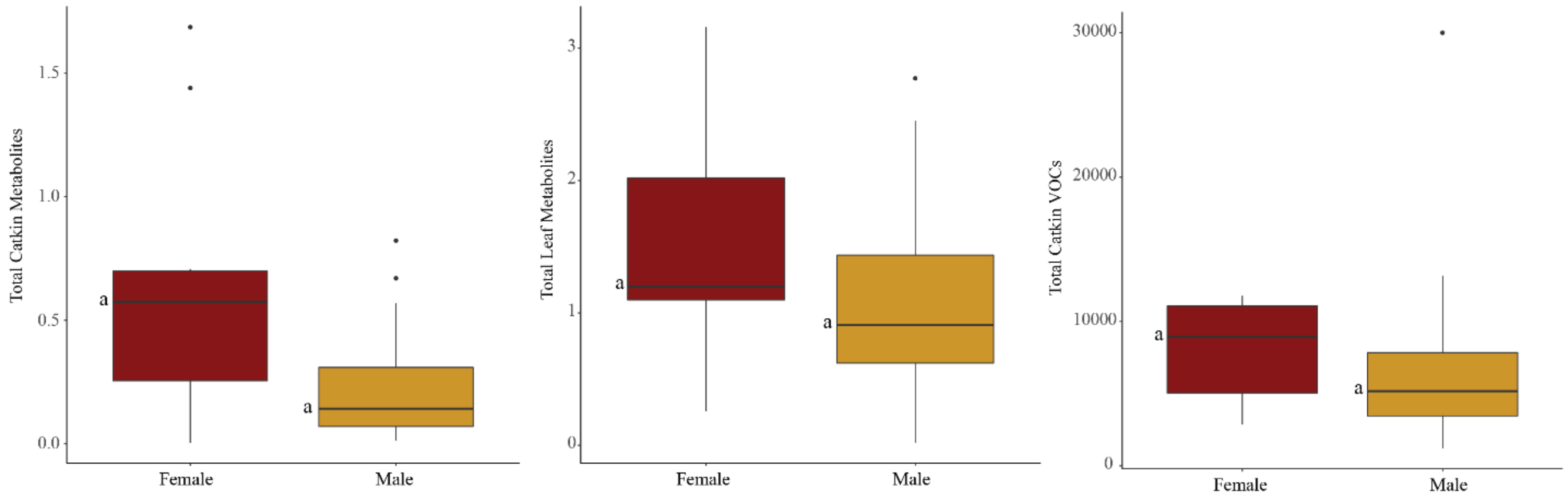


**Supplemental Figure 2.** Total concentration of catkin metabolites, leaf metabolites and catkin VOCs for male and female trees. Letters to the left of boxes indicate significantly different means (p-value < 0.05) determined by one-way ANOVA test.

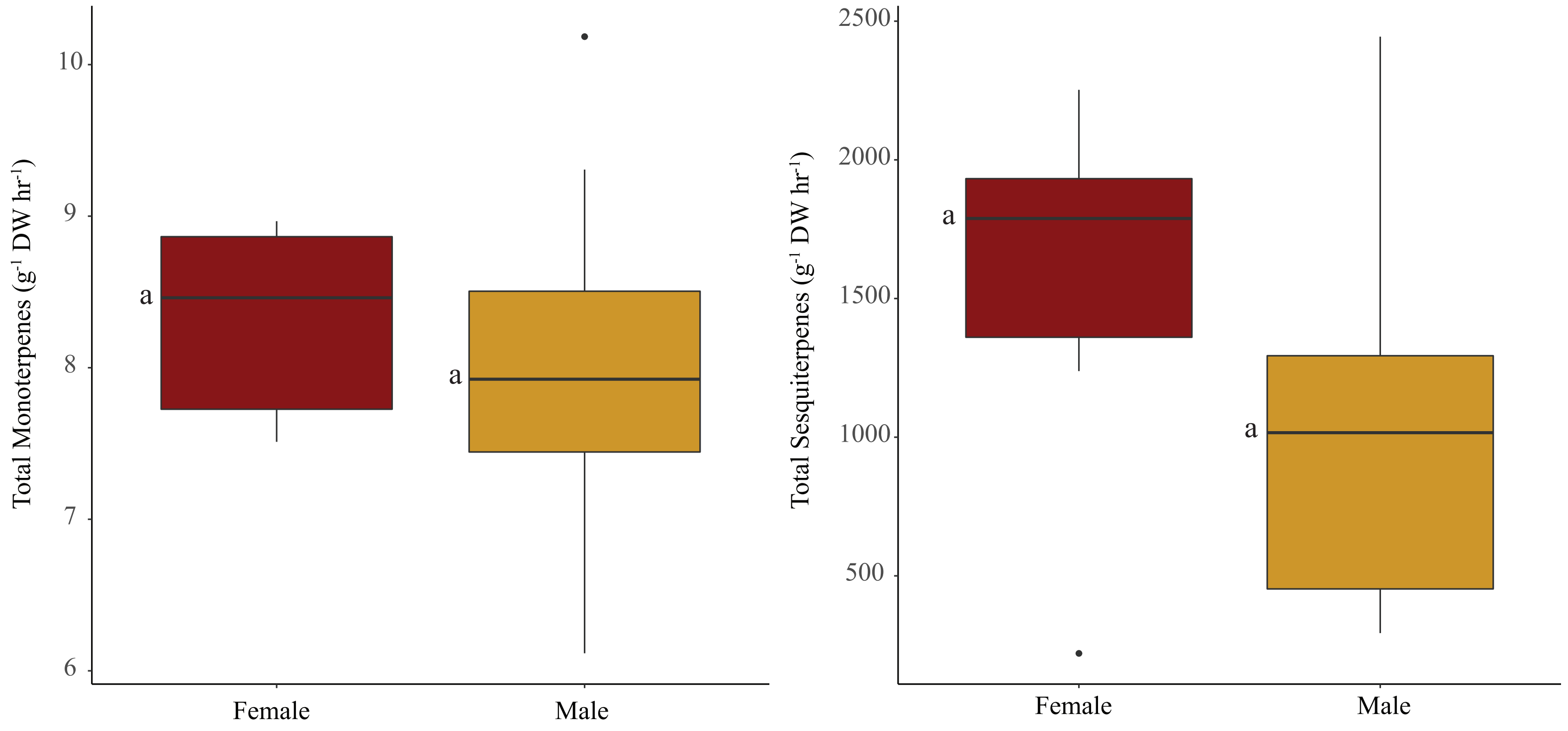


**Supplemental Figure 3.** Total concentration averages of monoterpenes and sesquiterpenes VOCs for male and female trees. Letters to the left of boxes indicate significantly different means (p-value < 0.05) determined by one-way ANOVA test.

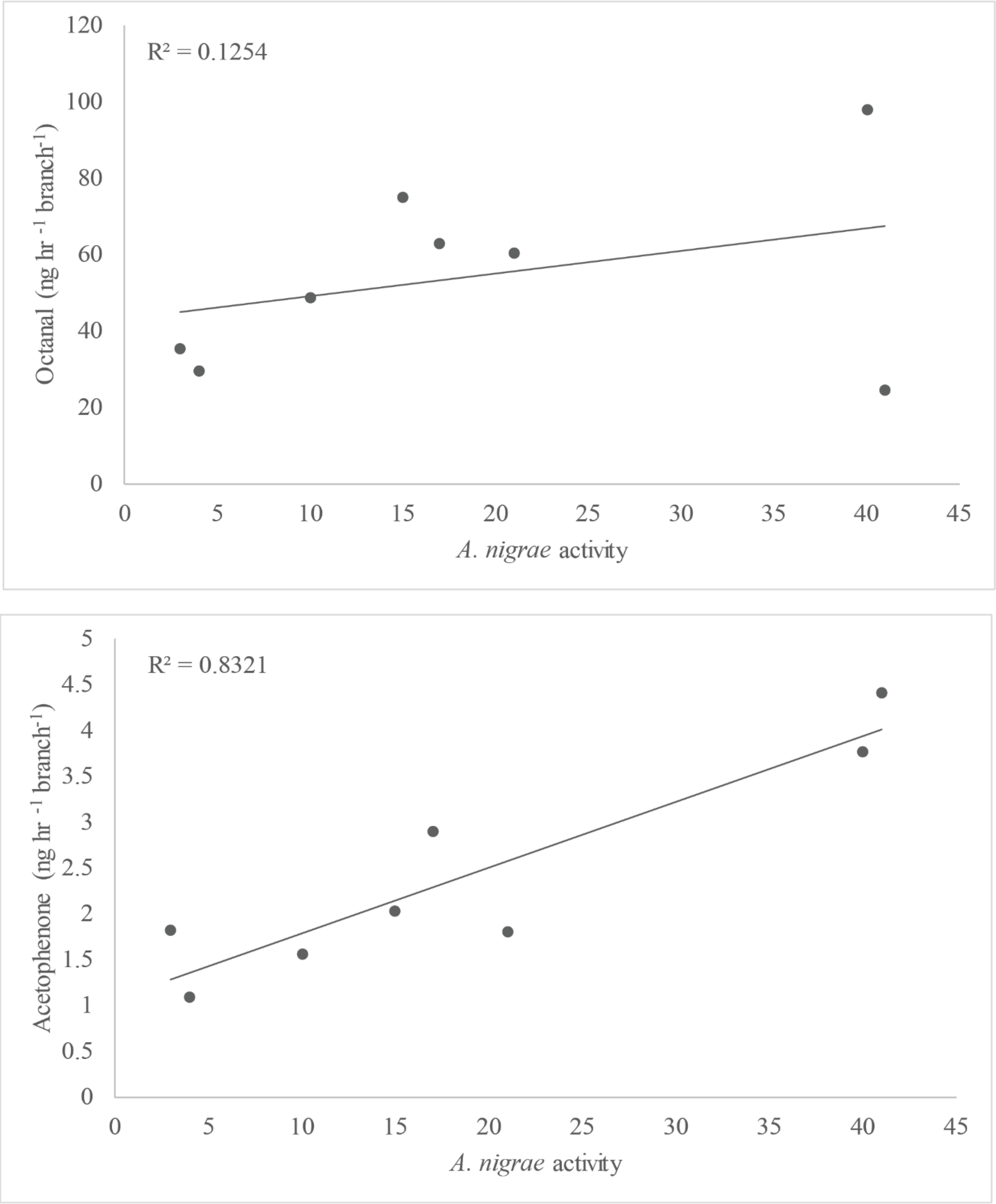


**Supplemental Figure 4.** Correlation of *A. nigrae* with VOC compounds.

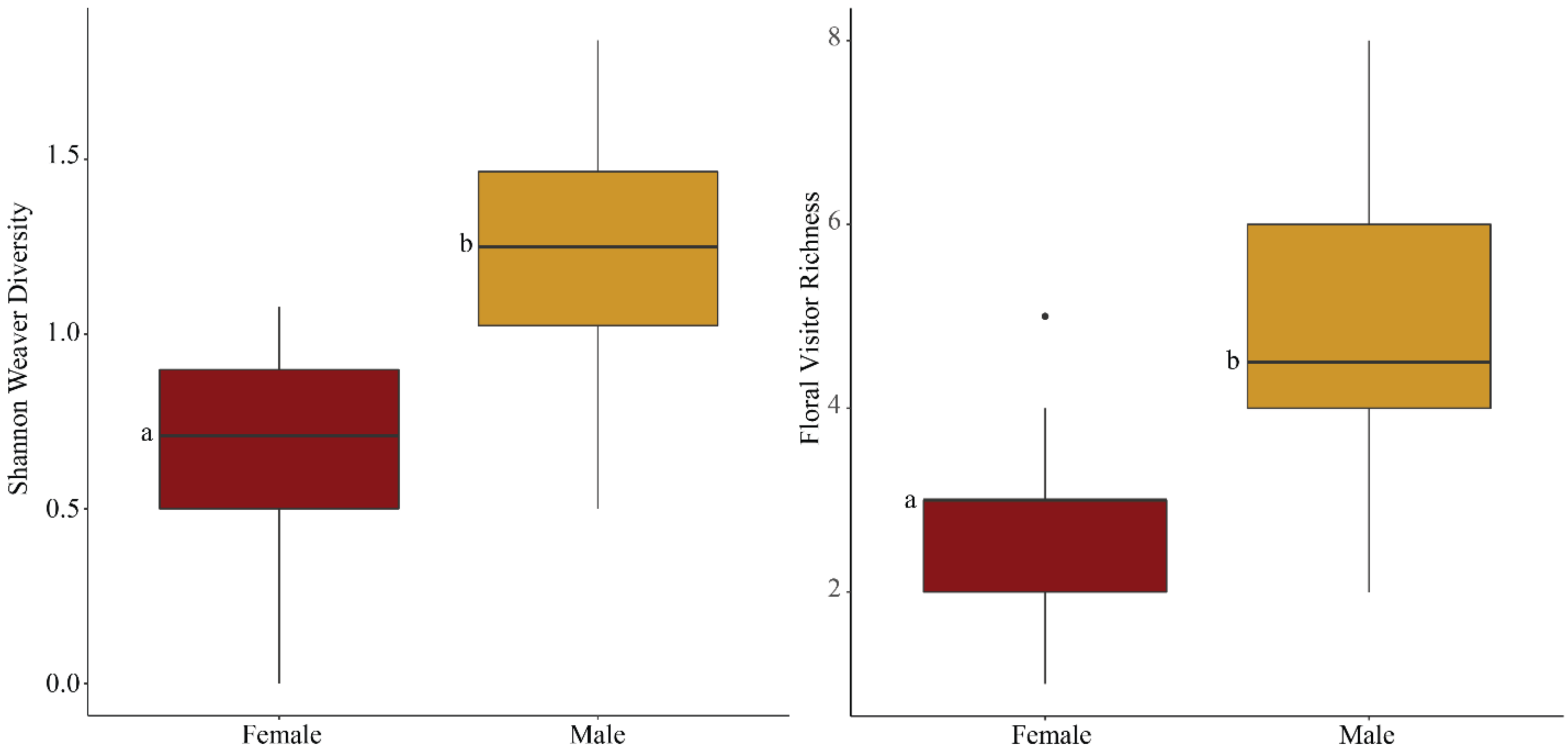


**Supplemental Figure 5.** Average values of species richness and Shannon-Weaver diversity for female and male trees from year analysis model. Letters to the left of boxes indicate significantly different means as determined by a Tukey's HSD (p-value < 0.05) for each separate nested ANCOVA model.
